# Expression and In-silico Structural analysis of N-acetylglucosamine transporter, Ngt1 in model fungal pathogen, *Candida albicans*

**DOI:** 10.1101/2025.04.14.648653

**Authors:** S. Haseena, Nadimpally Sai Tharun Goud, Soumita Paul, Swagata Ghosh, Sarika Sharma, Kongara Hanumantha Rao

## Abstract

**Introduction:** The commensal yeast, *Candida albicans*, has the enormous metabolic flexibility to utilize alternative carbon sources such as N-acetylglucosamine (GlcNAc) at the infection sites, thereby facilitating its adaptive strategies to colonize and thrive within diverse host niches. In *Candia*, GlcNAc specific transporter (Ngt1) imports GlcNAc at cell surface to induce signaling through the master regulator Ngs1 at the chromatin level. Thus, GlcNAc signaling is responsible for changing its habitat from commensal to pathogenic life style.

**Objective:** The current study aims at understanding the expression, structural and functional intricacies of Ngt1 that determines to GlcNAc sensitivity, specificity and its transport which is a prerogatory step in the GlcNAc signaling process in Candida.

**Method:** The 3-D structure of Ngt1 was predicted using AlphaFold. We have carried out docking studies using AutoDoc suite for native and Site directed mutant versions of Ngt1 with GlcNAc to determine role of critical amino acids and further, tunnel analysis to reveal the transport mechanism. These findings lead to the designing structure-based Ngt1blockers via virtual screening of natural compound libraries.

**Results:** Our docking studies on Ngt1 with GlcNAc revealed the useful insights in to mechanism and the role of critical amino acids (Ala 324, Ile 325, Tyr 329, and Asp 332) responsible for interaction. In our analysis, Asn 195, located within transmembrane helix and part of the UNC-93-like transmembrane regulatory protein domain, was identified as critical residue for GlcNAc interaction compared with other mutated residues. Interestingly, the tunnel analysis of Ngt1 transporter revealed presence potential pathways in Ngt1 protein for effective GlcNAc transport.

**Conclusion:** This study provides basic insights in to the structural aspects of Ngt1 and its interaction with GlcNAc that can be further explored for the development effective therapeutic agents for control of infection caused by the Candida.

Graphical Abstract:
Structural and functional analysis of Ngt1 protein.
Flow chart depicting Ngt1 3D modeling (AlphaFold2), docking studies Ngt1-wild type and SDM proteins with GlcNAc ligand, tunnel analysis reflecting transport of GlcNAc, and virtual screening to identify potential inhibitors (Brucine).

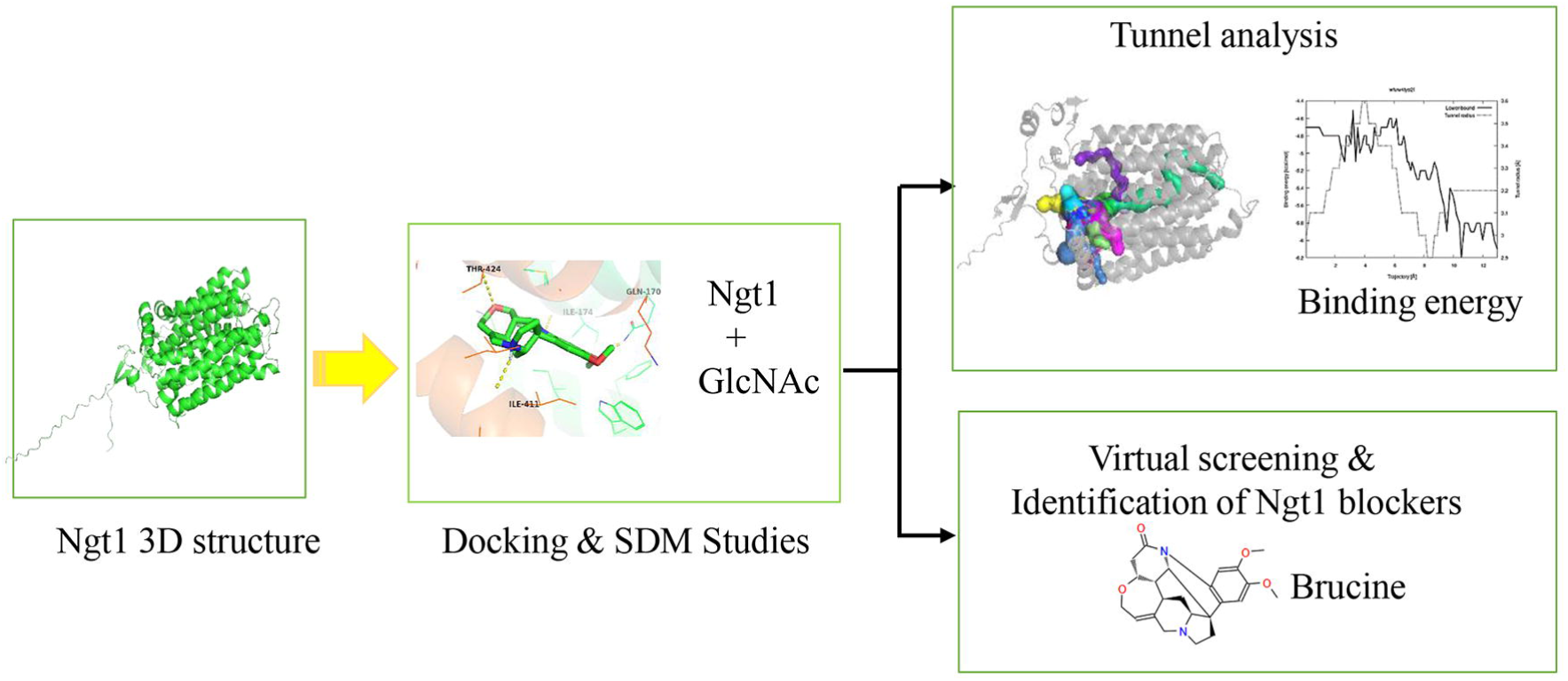

## INTRODUCTION

*Candida albicans*, while being a commensal part of the human microbiota, is an opportunistic pathogens that can lead to both superficial and serious infections, particularly in individuals with weakened immune system [1, 2, 3]. The increase in immunocompromised individuals, due to factors like radiotherapy, chemotherapy, organ transplants, and AIDS has led to a rise in Candida infections [4, 5, 6]. The morphological transition facilitates virulence and infection as the small budding cells are better suited to disseminate in the bloodstream and long hyphal filaments promote invasion into tissues and biofilm formation [7, 8, 9]. In the human host, Candida exists in different morphogenetic forms like yeast, hyphae, peudohyphae, white, opaque etc., [9, 10]. These morphogenetic transitions are induced by diverse factors like; elevated temperature (37◦C), neutral or alkaline pH of the ambient medium, CO2, serum, interaction with a solid extracellular matrix, and certain nutrients including GlcNAc and amino acids [11, 12, 13]. However, GlcNAc stands out as being one of the strongest inducers of hyphal growth.

GlcNAc available from the host tissue enters in to the Candida cells through GlcNAc specific transporter, N-acetylglucosamine transporter (Ngt1) at cell surface [14, 15, 16]. The internalized GlcNAc, utilized in the metabolism to provide energy, carbon and nitrogen source [17]. and further, this GlcNAc translocates to the nucleus where it interacts and activates the master regulator Ngs1 (N-acetylglucosamine sensor) at chromatin level to induce genes involved in various cellular processes including GlcNAc transport (*NGT1*), GlcNAc metabolism (*HXK1, NAG1, DAC1* etc.), GlcNAc scavenging (*HEX1*), morphogenesis (*HWP1, ECE1* etc.) and other uncharacterized functions [18, 19, 20]. Thus, GlcNAc import at cell surface through Ngt1 transporter is prerequisite and critical step in GlcNAc signalling [21, 22] process (Figure 1).

**Figure 1.**
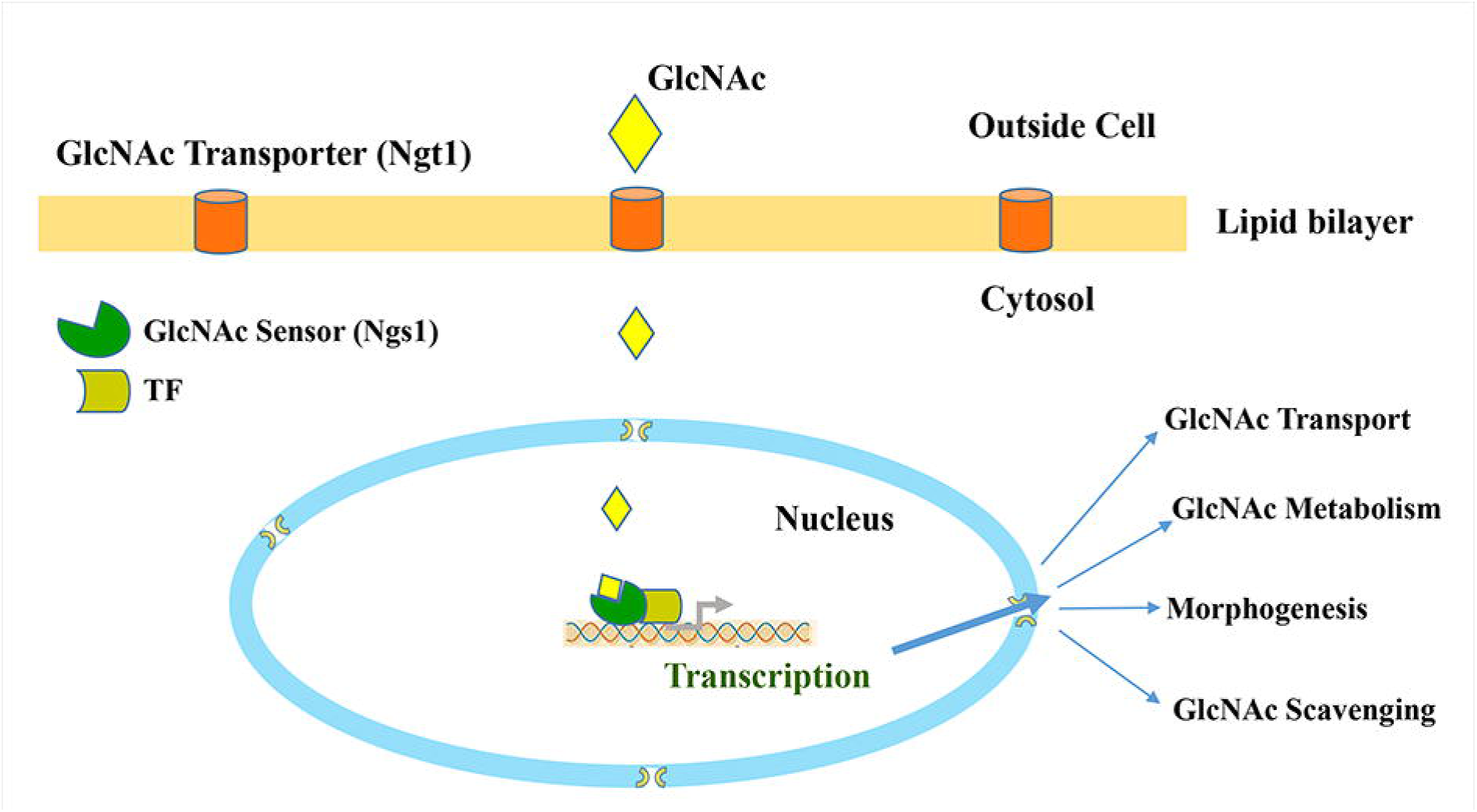
Model for GlcNAc import and its signaling. Exogenous GlcNAc is imported in to cytosol through GlcNAc specific Transporter, Ngt1 at cell surface. In the nucleus GlcNAc binds to GlcNAc sensor, Ngs1 to activate and transduce GlcNAc signaling. Ngs1 recruited to promoters of target genes with the help of Transcription factors like Rep1 [19, 20].

Although dynamics of Ngt1 in terms of its stability and protein levels at cell surface has been studied to some extent in our previous study [16], the structural insights that determine the function in terms of its GlcNAc binding properties and its transport mechanism are completely unknown. X-ray crystallography, Cryo electronmicroscopy (CryoEM) and NMR are commonly used methods for 3-D structural determination of proteins where the steps involved in protein expression, purification and crystallization very tedious and that might requires months to years of painstaking efforts to determine a single protein structure. Under such circumstances, in-silico analyses promise effective alternative approach for 3-D structural determination and there by understanding the functional dynamics of these membrane proteins [23].

A recently developed AlphaFold Protein structural Database [24] made it possible the prediction of the highly accurate protein 3-D structure from its amino acid sequence. AlphaFold is a novel neural-network based approach that incorporates protein biophysics with deep learning algorithms facilitating Protein structure predictions with accuracy at atomic level [25–27].

In the current study we have carried out computational analysis of the 3-D structure of Ngt1 and determine the specificity of its interaction GlcNAc ligand and further the transporter mechanism of Ngt1 that plays a dynamic role in GlcNAc import at the plasma membrane facilitating signaling mechanisms. Initially, the 3-D structure has been predicted by AplhaFold, a state of art tool. The binding interactions between Ngt1 and GlcNAc, the critical amino acids that make the compatible grooves in Ngt1 determined by molecular docking study using AutoDock [28]. The tunnel analysis revealed the identification of 13 tunnels and Tunnels 1 and 12 form the potential channel for effective transport of GlcNAc. Further, with the aim to identify the blockers for Ngt1 that could inhibit the GlcNAc binding, we have carried out virtual screening by implementing Rapid Pharmacophore-Based Screening (RPBS) platform, the MTiOpenScreen tool [29] and identified three compounds (ZINC000043552589, ZINC000044404209 and ZINC000005512974) as potential blockers with score around −11.5 to −10.1 (with number of interaction range from 8 to 22).

## MATERIALS AND METHODS

### Methodology and materials used in wet lab experiments

#### Candida Culture conditions

YPD (1% Yeast Extract, 2% Peptone, 2% Glucose, w/v) broth and agar (2 % agar) were used for regular culture of *C. albicans* strains at 30LJ incubator. For induction purposes, overnight-grown saturated cultures were added to the SD media for the growth of the culture until the OD reached 0.6. Then the cells were washed twice and re-suspended in YNB media containing different carbon sources (glucose, GlcNAc). Cells were collected after 90 minutes of induction and proceeded to the downstream experiments

#### Immunoblotting

Immunoblotting was performed as described by Rao et. al. (Hanumantha Rao et. al., 2022). Protein was isolated from the *C. albicans* cells using ice-cold TES lysis buffer having PhosSTOP and Complete EDTA-free protease inhibitor from Roche. For doing Western Blot analysis, 20 μg- 20 ng of proteins from gels were electrotransferred to Hybond C Extra membrane (Amersham Biosciences)(Lenoir et. al., 2009). The blot was blocked using 3% skimmed milk in a buffer containing 10 mM Tris, pH 7.4, 15 mM NaCl, 0.05% Tween-20. For detection of GFP signals, membranes were incubated with an anti-GFP polyclonal antibody (Abcam, cat no. ab290) at a dilution of 1:15000 followed by a peroxidaseconjugated secondary antibody (Amersham Biosciences, UK) at a dilution of 1:20000 (Hanumantha Rao et. al., 2022). An enhanced chemiluminescence (ECL prime) detection system (Advansta, USA) was used for antibody detection according to the manufacturer’s instructions. Chemiluminescence signals were developed in the Kodak Carestream x-ray film.

#### Microscopy

Fluorescence images were captured by using a a Nikon 80i inverted microscope equipped with a Nikon digital DXM1200C camera a with the excitation wavelength 440-475 nm for GFP fluorescence. Images were processed by using Adobe Photoshop version 5.5 (Adobe Systems, Corp., San Jose, CA) for necessary adjustment.

### Methodology used in In-silico analysis for the Study of Ngt1 in Candida albicans Molecular Docking: Methodology

#### Retrieval of Ngt1 Structure

The importance of using publicly available biological data sources, such the Candida Genome Database, has been well recognized [30]. Protein structural and functional characterisation relies heavily on the retrieval of amino acid sequences and structural information [25]. In the case of the Ngt1 transporter, the amino acid sequence was retrieved from the Candida Genome Database and subsequently submitted to the AlphaFold server for high-confidence structural prediction [25, 26]. The Ngt1 transporter protein’s three-dimensional structure was computationally predicted using AlphaFold server, a cutting-edge AI-based platform for making precise protein folding predictions. The Structural Analysis and Verification Server (SAVES) were used to validate the initial model, which had 509 residues. The protein structure’s stereochemical quality was assessed using PROCHECK analysis, which mainly used Ramachandran plot analysis to focus on side chain and backbone geometry, and the ERRAT study [37], which showed the model’s remarkable non-bonded atomic interactions and general stereochemical dependability [38]. The secondary structure of the protein was analyzed, revealing a complex arrangement of alpha-helices, beta-sheets, and turns through CFSSP: Chou and Fasman Secondary Structure Prediction Server [39].

#### Docking study

The ligand, N-acetylglucosamine, was obtained from the PubChem database [33], whereas the structure of the Ngt1 transporter receptor was retrieved from the AlphaFold Protein Structure Database in PDB format. Hydrogen atoms were added, rotatable bonds were defined, and partial charges were assigned using Gasteiger charges for the ligand [28]. In order to ensure conformational flexibility and reliable binding affinity prediction during simulations, the receptor was prepared by assigning Kollman charges, merging polar hydrogens, and assigning AD4 atom types [39]. The binding site was encapsulated in a grid box with dimensions of 126 × 74 × 126 points along the X, Y, and Z axes and a grid spacing of 0.375 Å. The docking grid, centered at coordinates (−9.617, −5.605, 14.582), utilized receptor atom types (A, C, H, HD, N, OA, SA) and ligand atom types (A, C, HD, N, NA, OA, SA), with affinity maps and an electrostatic potential map generated to enable precise ligand-receptor interaction mapping. Energy fields and spatial arrangements for docking were established using AutoGrid to create grid maps. After that, AutoDock was then executed to predict binding poses and affinities by sampling and scoring ligand conformations efficiently.

#### Site-Directed Mutagenesis

To elucidate the functional role of specific residues in the Ngt1 transporter, site-directed mutagenesis [40] was performed at three key positions: 80(Asparagine), 195(Asparagine), and 320(Glutamine). Residues 80, 195, and 320 were chosen for mutation because of their participation in conserved and significant functional regions found by InterProScan analysis [41]. The MFS (Major Facilitator Superfamily) domain, which is critical to residue 80 and is designated by Gene3D (G3DSA:1.20.1250.20) and SUPERFAMILY (SSF103473), is essential for the transport of substrates across membranes, indicating that it plays a role in transport-related functions. Residue 195 falls within a predicted transmembrane helix, as indicated by TMHMM and Phobius [42], and is also part of the UNC-93-like transmembrane regulatory protein domain (Pfam: PF05978, IPR051617) [43].Transmembrane helices are essential for substrate recognition and movement across the membrane. Lastly, residue 320 is located in another transmembrane domain predicted by TMHMM and Phobius [42], further highlighting its potential role in forming the transport channel or mediating substrate interactions. These findings suggest that the selected residues are conserved and play pivotal roles in the protein’s structural integrity and functional mechanisms, making them prime candidates for mutagenesis studies to elucidate their precise contributions to transport activity. The mutagenesis process involved the following steps.

The Chimera software was employed to make targeted mutations [44]. The original positions were mutated to alanine, a process that simplifies the side-chain interactions and allows for the assessment of the importance of each residue’s contribution to ligand binding. The modified sequences were then used to create pdb files for each mutant variant, which were vital for further computational docking and analysis. Following the mutagenesis, energy minimization was implemented to ensure that the mutant structures were in stable conformations Utilizing PyMOL [45] and PDBViewer [46]. The aim was to remove any steric clashes and optimize the geometries of the mutants before further studies. The docking procedure was repeated for each of the mutant variants to compare the binding affinities and interactions with N-acetylglucosamine.

The tunnel analysis of the Ngt1 transporter protein was performed using the Caver Web Server [47] to identify and characterize potential pathways for substrate transport. The Caver Web Server analyses protein tunnels by identifying probable paths that connect the protein’s interior to its surroundings environment. By mapping the solvent-accessible voids in the protein, it computes tunnels using a method based on a Voronoi diagram. Tunnel qualities were assessed by computing parameters like length (distance from the active site to the protein surface), curvature (path deviation, impacting substrate movement), throughput (an integrative efficiency metric combining the other parameters), and bottleneck radius (narrowest point determining substrate size compatibility). The technique uses static protein structures or pre-calculated molecular dynamics as input and incorporates clustering of pathways according to their geometric similarities [48]. The algorithm creates accurate tunnel paths for functional investigation by taking into account solvent accessibility, probe size, and atomic radii.

#### Virtual screening

begins by converting the obtained structural data into a machine-readable format suitable for computational analysis. The docking results obtained Auto-Dock served as inputs for the RPBS, which allowed for the identification of compounds that could serve as likely blockers of Ngt1 activity. The RPBS platform offers the MTiOpenScreen tool [20], an advanced bioinformatics resource designed for virtual screening and molecular docking. It integrates a variety of docking algorithms, including AutoDock for robust ligand-receptor binding predictions using Lamarckian Genetic Algorithms, and AutoDock Vina, which enhances speed and accuracy with advanced scoring functions. The tool also supports flexible docking to account for ligand conformational variability, rigid docking for efficient initial screenings, and grid-based scoring algorithms to evaluate binding affinities with spatial interaction potentials. Users can screen compound libraries, such as natural products, for potential inhibitors or modulators. These customizable settings, including adjustable docking runs, make MTiOpenScreen a precise and versatile tool for drug discovery research. This approach identified several compounds from the NP database [39] that could potentially block Ngt1 activity.

## RESULTS AND DISCUSSION

### Ngt1-GFP expression is highly sensitive to even very low concentration of GlcNAc

N-acetylglucosamine transporter, Ngt1 is GlcNAc specific transporter involved in the import of GlcNAc at cell surface [14]. GlcNAc entry through Ngt1 is a prerogatory step in GlcNAc signaling and metabolism [19, 20]. Considering the critical role of GlcNAc in *C. albicans* virulence, it has become imperative to understand expression kinetics and structural dynamics of Ngt1 that can determine Ngt1 levels at cell surface and its activity respectively.

In the current research, to study the expression kinetics of Ngt1, we have used a strain expressing a GFP fusion protein (Ngt1-GFPLJ) which was used in our previous work [16]. To determine the sensitivity of Ngt1-GFP expression i.e the minimum concentration of GlcNAc that can induce Ngt1 expression, we have exposed the Candida cells to the decreasing concentrations of GlcNAc starting from 2% (90 mM) at the rate of 6 fold serial dilation for 2 hrs as shown in the figure 2. We have noticed Ngt1-GFP localization signals till 1.96 µM GlcNAc concentration (Figure 2A). Simultaneously, we have carried out western blotting analysis to detect the Ngt1-GFP signals by using anti-GFP antibody for crude lysates obtained from the cells induced for 2 hrs with 2.5 mM GlcNAc which is optimum concentration used for GlcNAc induction experiments [16, 20]. In the immunoblot (Figure 2B) performed with different concentrations of protein isolated from 120 min time point collected cells, signals could be detected even with 20 ng protein loaded that lent further credence to the prolific and sensitive nature of the protein

**Figure 2.**
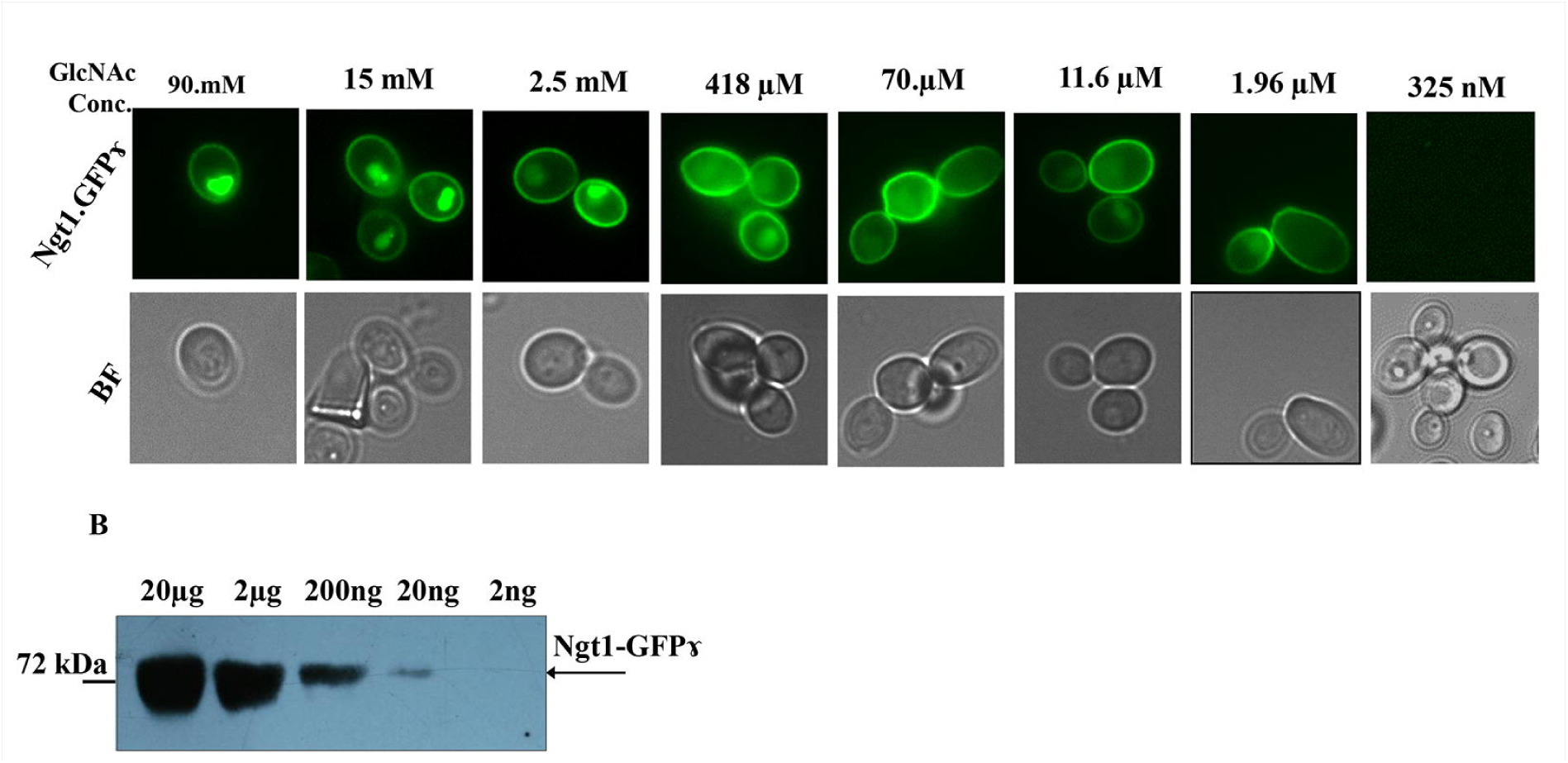
Ngt1-GFP expression is highly sensitive to GlcNAc. A. Ngt1 is expressed even at 2µM GlcNAc concentration. Localization pattern of Ngt1-GFPLJ at variable concentrations of GlcNAc with in the range of 90 mM to 325 nM. *C. albicans* strain expressing Ngt1-GFPLJ were grown till exponential phase in YPD and washed twice in SC basal medium induced in GlcNAc (SC-GlcNAc) of indicated concentrations i.e from 90 mM to 325nM (6 fold serial dilutions). They were allowed to grow for 2hrs and observed under fluorescence microscope. **B. Ngt1-GFP expression is highly prolific.** Western blot indicating the expression of Ngt1-GFPLJ at 90 min after addition of GlcNAc (2.5mM). Different amounts of total protein, 20µg to 2ng was loaded on 12% SDS-PAGE gel and processed for western blotting analysis using anti-GFP antibody.

#### 3-D structural prediction for N-acetylglucosamine transporter, Ngt1 and ligand GlcNAc

Ngt1 protein sequence was retrieved from Candida Genome Database [30]. The protein, length of 509 amino acids containing 12 transmembrane-spanning domains, belongs to major facilitator superfamily [31, 32]. From primary sequence, we have predicted the 3-D structure of Ngt1 by using AlphaFold2 web tool [24] (Figure 3). The resulting Ngt1 structure were processed and formatted appropriately for subsequent computational analyses (described in methods). Similarly, the 3D structure of the ligand N-acetylglucosamine in SDF format is obtained from the PubChem [33].

**FIGURE 3:**
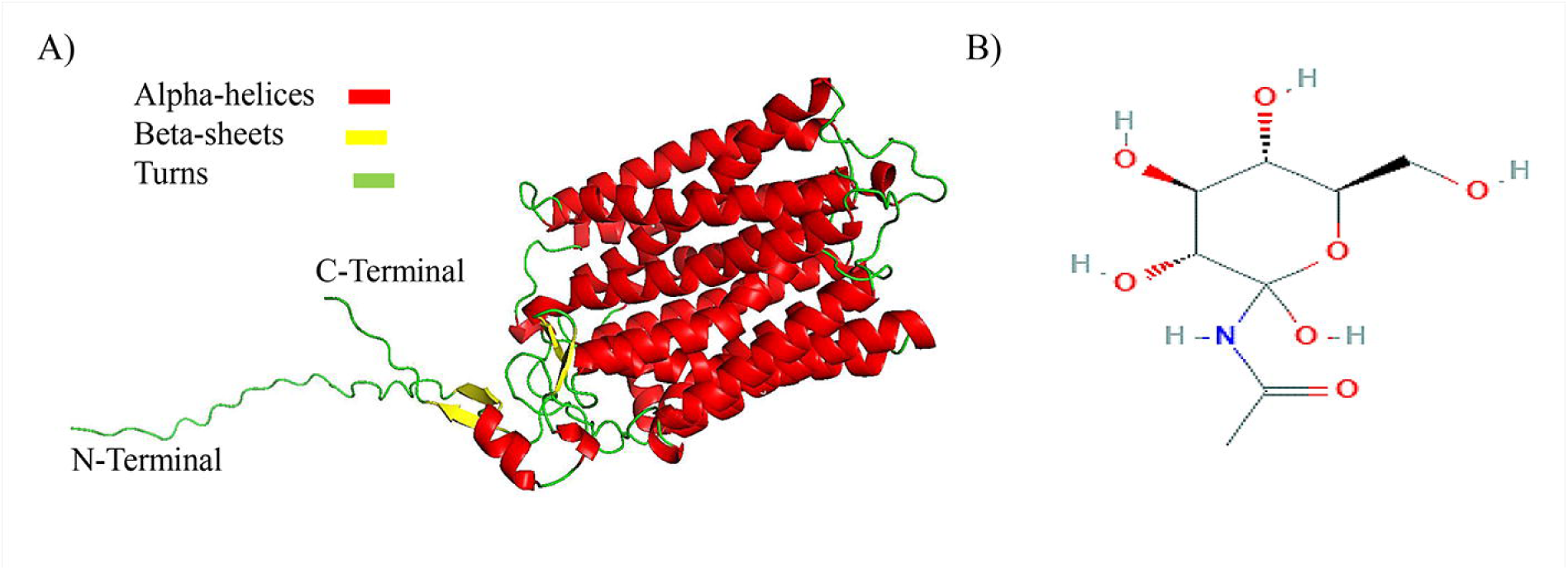
The 3-D structure details GlcNAc transporter, Ngt1 and its ligand, GlcNAc. A) GlcNAc transporter protein 3-D structure was predicted using AlphaFold tool that provides high accuracy of structure predictions. B) The 2D structure of the ligand N-acetylglucosamine is obtained from the pubchem.

#### Validation of Ngt1 3-D model

The initial model obtained from AlphaFold (Figure 3A) further validated by using tools like SAVES and ERRAT as described in Methods section. An overall quality factor of 98.27 % was obtained from the ERRAT study which shows there are minimal non-bonded atomic interactions. Subsequently, PROCHECK analysis revealed that 8.0% of residues were discovered in the Ramachandran plot’s additional allowed regions, whereas 92.0% were identified in the most favoured regions. Importantly, the model’s great structural accuracy was demonstrated by the absence of residues in the generously allowed or disallowed regions. These findings satisfy the requirements for high-resolution structures, which are often defined by more than 90% of residues in the most favored regions. The secondary structure of the Ngt1 transporter consists of 509 residues, with 356 residues (69.94%) adopting an alpha-helix conformation, indicating a predominantly helical structure. Additionally, 18 residues (3.54%) form beta-sheets, contributing a smaller portion of beta-strands to the protein’s framework. A significant portion, comprising 135 residues (26.52%), is involved in coils (turns/loops), providing flexibility and connectivity between the structural elements This distribution demonstrates the protein’s complex folding and structural stability, which makes it a strong candidate for additional functional and interaction study.

Here (Figure 3), we present a structure of N-acetylglucosamine transporter, Ngt1 generated using AlphaFold. The transporter shown embedded in a plasma membrane and N-terminal Glu, Lys, Asp, Gln, Thr, Ile, Asn, Ser, Leu, Val and Pro residues) and C-terminal (Lys, Ser, Thr, His And Glu residues) are protruded in to cytosol and that are rich hydrophilic amino acids. Very recently, for a few membrane transporters, 3-structures have been solved by XRD and CryoEM. In these studies, AlphaFold predicted models are shown to be quite accurate with experimental derived structures [23, 34]. Further, these AlphaFold predicted 3D models were successfully used for simulation studies.

#### Docking studies of Ngt1 with GlcNAc: Docking studies with AutoDoc suite revealed critical amino acids of Ngt1 for GlcNAc specificity fi

Docking studies have been conducted for Ngt1 (wild type and Site directed mutagenesis-SDM proteins) with GlcNAc to determine the binding affinities, interaction patterns, and potential inhibitors of the Ngt1 transporter. PyMol was used to visualize the interactions between the GlcNAc and the target protein Ngt1 transporter. Table S1 shows the Ngt1 amino acids that interact with the GlcNAc. Figure 4A and 4B shows the 2-D and 3-D pictures of the Ngt1 docked GlcNAc complexes illustrating the interactions.

**FIGURE 4:**
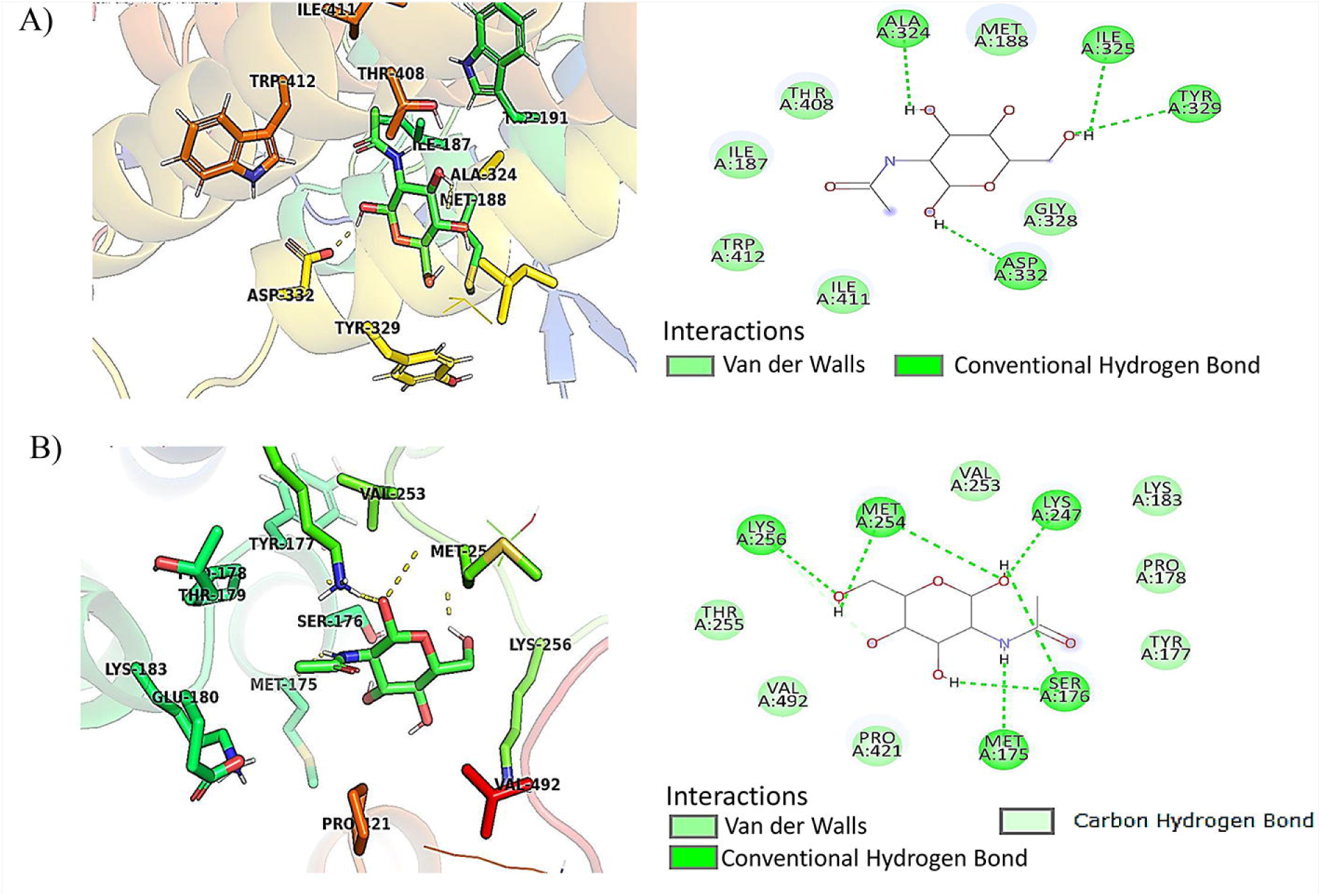
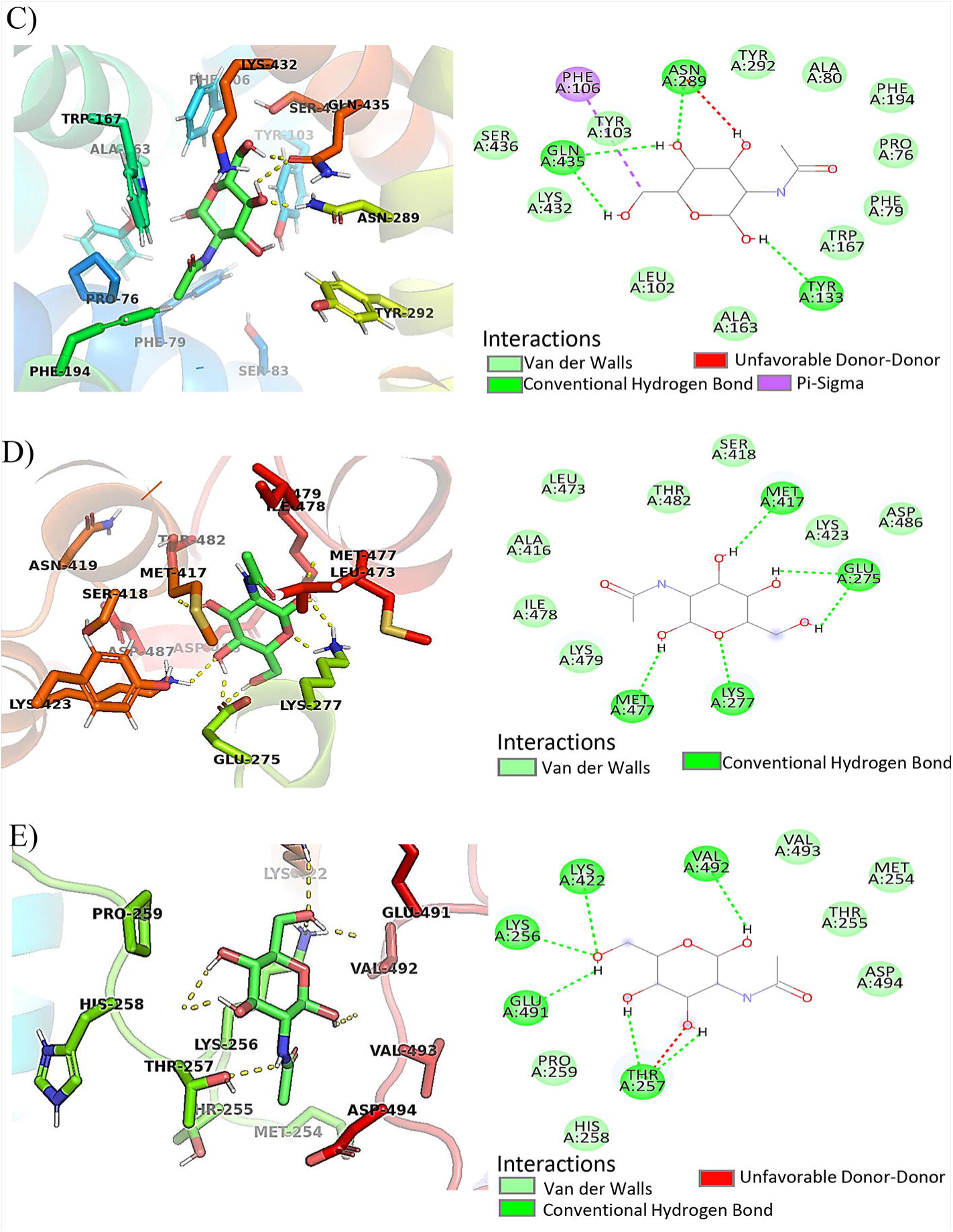
3-D Structure interactions of GlcNAc transporter, Ngt1 and its ligand, GlcNAc. **A) Native 3D and 2D interactions with 4 H-bonds are represented (**Ala 324, Ile 325, Tyr 329, And Asp 332)**. B) Mutation at three residues** (80, 195, and 320) 3D and 2D interaction with 5 H-bonds C) Mutation at 80 residue Asparagine 3D and 2D with 3 H-bonds D) Mutation at 195 residue Asparagine 3D and 2D with 4 H-bonds E) Mutation at 320 residue Methionine 3D and 2D interaction with 4 H-bonds.

The docking results highlight the impact of mutations on the binding affinity of GlcNAc with the Ngt1 membrane protein and its variants. The native Ngt1 has a binding energy of −5.4 kcal/mol, primarily stabilized by hydrogen bonds with residues ALA 324, ILE 325, TYR 329, and ASP 332, alongside hydrophobic interactions involving VAL 122, LYS 123, and ILE 242 (Figure 4A). The triple mutant (Mutations at 80, 195, and 320) (Figure 4B) improves the binding energy to −5.7 kcal/mol, likely due to additional stabilizing hydrogen bonds with SER 176, LYS 247 and MET 254 (Table S1). Mutation 195 demonstrates the highest binding affinity with an energy of −6.2 kcal/mol, supported by interactions with ASN 80, TRP 167, TYR 292, GLN 320 and GLN 435, probably indicating its critical role in ligand stabilization. Mutation 80, on the other hand, exhibits the weakest binding affinity (−5.0 kcal/mol), with fewer hydrogen bonds (TYR 113, ASN 289, GLN 435) and limited hydrophobic interactions. Mutation 320 shows a moderate binding energy of −5.7 kcal/mol, with key interactions involving LYS 256, LYS 422, and THR 257. These observations suggest that specific residues, particularly at position 195, significantly contribute to ligand binding in the membrane environment. These findings align with previous studies that emphasize the role of hydrogen bonding and hydrophobic interactions in ligand recognition and stabilization in membrane protein systems [35] (Tables S1, S2 and S3).

#### Ngt1 tunnel analysis to identify potential pathway for GlcNAc transport

The tunnel analysis of the Ngt1 transporter protein was performed using the Caver Web Server to identify and characterize potential pathways for substrate transport.

The tunnel analysis of the Ngt1 transporter protein was performed using the Caver Web Server to identify and characterize potential pathways for substrate transport (Figure 5). Four essential parameters like bottleneck radius, length, curvature, and throughput are used to evaluate tunnels (Table S4). Tunnel 1 was chosen as the most effective channel for substrate transport out of the 13 tunnels found because it had the optimal characteristics that largest bottleneck radius (2.5 Å), the shortest length (11.8 Å), the lowest curvature (1.3), and the maximum throughput (0.84). On the other hand, Tunnel 13 had the lowest throughput (0.04) due to its long length (66.2 Å), high curvature (2.4), and tight bottleneck radius (1.0 Å). Tunnel 1’s favorable features indicate that it is the Ngt1 transporter’s principal functional conduit, which corresponds to its physiological purpose.

**FIGURE 5:**
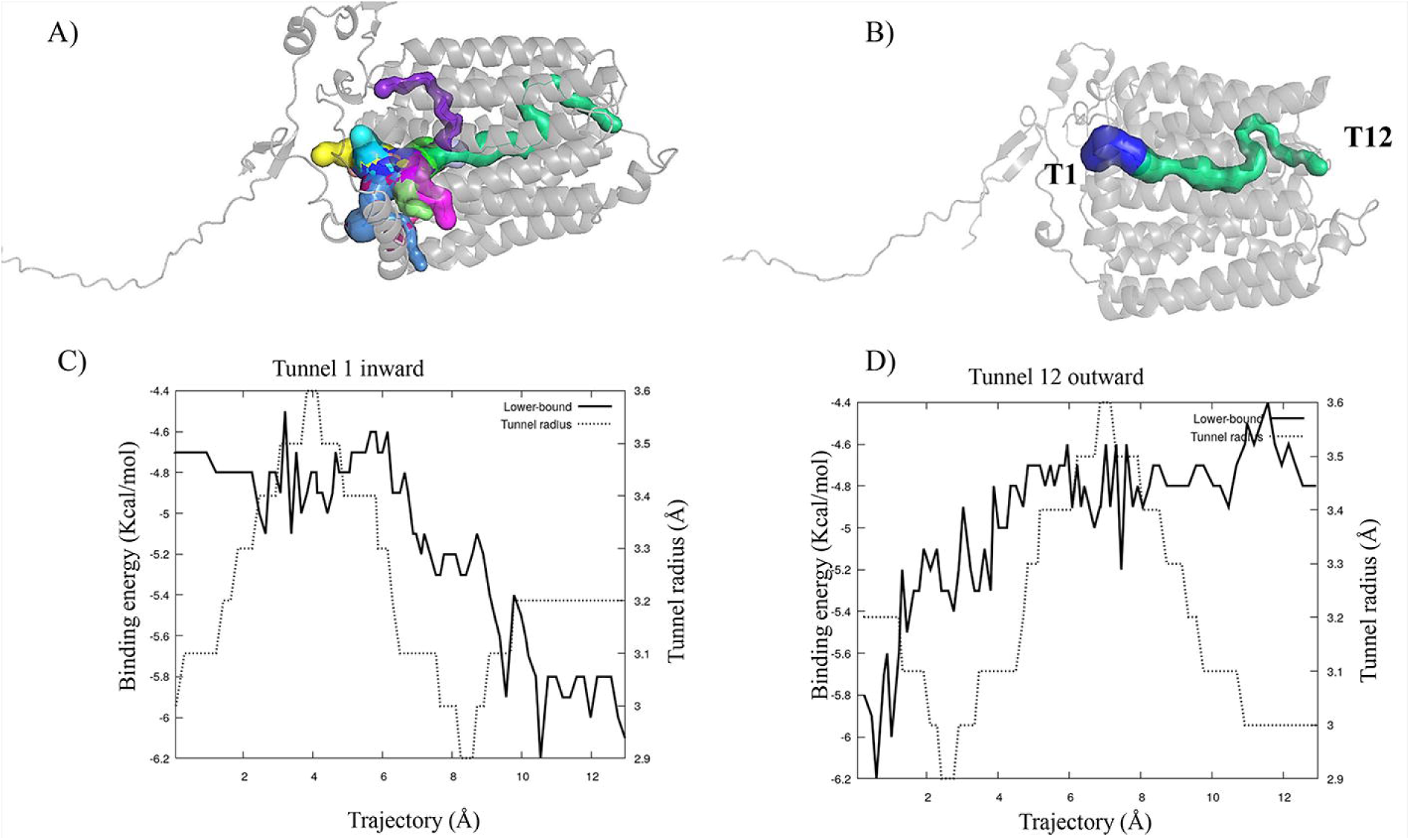
Ngt1 transporter tunnels, generated via the Caver Web Server. A) Total 13 tunnels were identified in the NGT1 transporter B) Tunnel 1 (T1-green) and Tunnel 12 (T12-dark blue) with high bottleneck and length. Energy landscape of the tunnel pathway associated with the GlcNAc transport, generated via the Caver Web Server. C) Energy profile of the tunnel 1 formed by the GlcNAc-binding site shows entry, and, D) Energy profile of the tunnel 12 formed by GlcNAc-binding site shows exit.

#### GlcNAc Binding Analysis in the identified tunnel

The CaverDock energy profile graphs show that ligands were successfully transported through interconnecting tunnels, notably Tunnels 1 and 12. The ligand’s inward movement is depicted by the first graph, (Figure 5C) which corresponds to Tunnel 12. The binding energy fluctuates as the ligand moves inward, beginning at about −5.4 kcal/mol close to the tunnel opening. Transient stabilization regions, where the ligand interacts with the tunnel walls, are shown by these fluctuations. When the ligand reaches a critical location, most likely the interface or a junction with Tunnel 1, the energy decreases significantly to −6.2 kcal/mol toward the end of the trajectory, indicating considerable stability. Concurrently, the tunnel’s radius shrinks, reaching a bottleneck at about 3.0 Å before stabilizing. This narrowing suggests that the ligand encounters structural constraints but adapts effectively to proceed along the tunnel path. As the ligand stabilizes at the junction or exit of Tunnel 12, the binding energy begins at about −4.6 kcal/mol in the inward tunnel 1 graph, which corresponds to Tunnel 12 and outward transport. The binding energy increases as the ligand travels outward, ranging from −4.8 to −5.0 kcal/mol. This increase suggests a gradual loss of stabilizing connections as the ligand passes through Tunnel 1 and gets closer to the protein’s surface. The tunnel radius varies along the route, including an earlier bottleneck (∼3.1 Å). This bottleneck supports the ligand’s successful outward transition via Tunnel 1’s structural constraints by matching the energy profile. The ligand is effectively transferred between both channels, as seen by the coupled energy profiles of Tunnels 12 and 1. As the ligand approaches a critical junction, the energy profile and radius variations confirm efficient ligand transport through the tunnels, with Tunnel 12 facilitating entry and Tunnel 1 enabling a smooth exit. Together, these profiles demonstrate the protein-ligand system’s structural and energetic flexibility by confirming efficient directed ligand transit between the two tunnels.

#### Virtual Screening for Ngt1 Blockers Using RPBS

To identify potential inhibitors of Ngt1, a Virtual Screening process was Implementation of Rapid Pharmacophore-Based Screening (RPBS) [36]. The top docking results revealed the following binding scores (Table S5 and S6): MolPort-001-742-110 (ZINC000043552589) showed the strongest binding with a score of −11.5 and 8 interactions, followed by MolPort-039-052-621 (ZINC000044404209) with a score of −11.3 and 9 interactions. MolPort-000-704-745(ZINC000005512974) had a binding score of −10.1 with 22 interactions with varying numbers of interactions. These results suggest that these compounds might effectively interfere with Ngt1 function. Here, we have chosen compound ZINC000005512974 that showed maximum number of interactions with optimum binding energy.

#### ZINC000005512974 Binding Analysis

The binding energy begins at −8.5 kcal/mol near the entry tunnel 12 and fluctuates before settling around −5.0 kcal/mol near the end of the trajectory. The blocker’s robust interaction with the tunnel, possibly occupying important binding sites along the channel, is indicated by this notable stabilization at lower energy values. The tunnel radius profile indicates narrow sections (∼3.0 Å) that match with regions of strong binding. This suggests that the blocker fits closely into the tunnel, physically obstructing the channel and limiting GlcNAc transport. Similarly, the binding energy varies during the trajectory in the outward of tunnel 1, beginning at −8.5 kcal/mol. The binding energy stays comparatively low, in contrast to a ligand that is freely transported, suggesting that the blocker is continuously stabilized inside the tunnel. The energy minima imply that the blocker creates stable contacts that are challenging to break, especially at constriction sites. The hypothesis that the blocker generates steric hindrance, rendering the channel inaccessible for GlcNAc, is further supported by the tunnel radius, which likewise exhibits significant narrowing (∼3.1 Å) Overall, the blocker’s energy profiles show that it successfully prevents GlcNAc from being transported through Tunnels 12 and 1. The tight binding and stable interactions, combined with the tunnel constriction, create a physical and energetic barrier that prevents the GlcNAc movement through the protein structure.

#### Comparative Analysis and Potential Blocking Mechanism

Since the blocker has a higher binding affinity than GlcNAc and has a different energy profile, it is possible that ZINC000005512974 functions as a competitive inhibitor (Figure 6). The transport of GlcNAc via the protein may be effectively constrained or delayed by the blocker’s occupation of the tunnel and higher binding affinity. This hypothesis is supported by the energy differences, where the blocker exhibits stronger interactions (lower energy values) that might prevent GlcNAc from successfully traversing the binding site. Overall, these findings point to a competitive relationship in which the blocker may sterically impede or modify the protein’s accessibility for GlcNAc binding and transport. Further structural and dynamic simulations would be useful in confirming this connection and its mechanistic impact.

**Figure 6:**
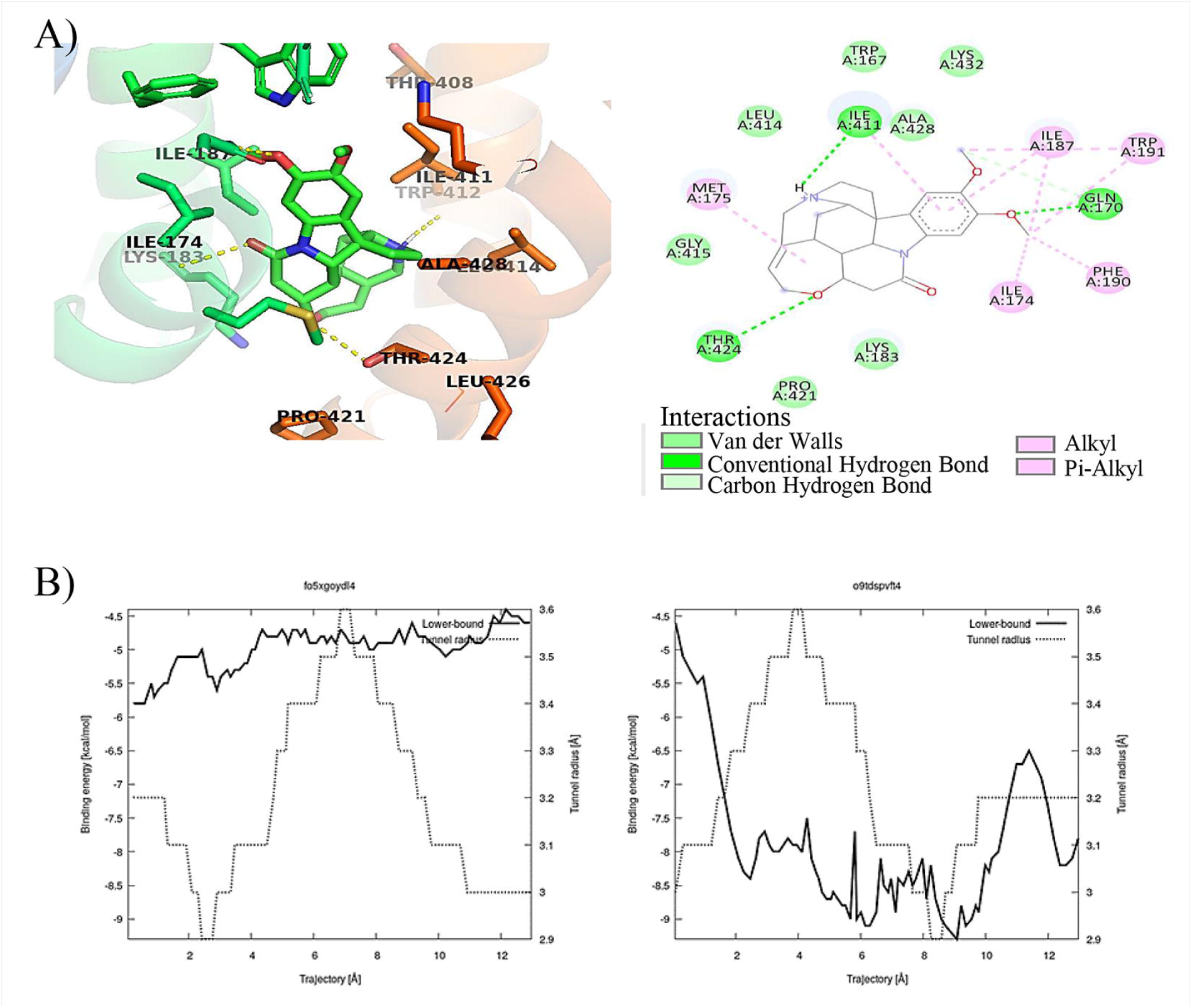
A) 3-D Structure interactions of Blockers. Ngt1 and its ligand ZINC000005512974 3D and 2D with 3 H-bonds. B) Energy landscape of the tunnel pathway associated with the blocker, generated via the Caver Web Server. Energy profile of the tunnel formed by the Blocker ZINC000005512974 binding site.

## CONCLUSIONS AND FUTURE PERSPECTIVES

The current study is novel on its own as it provides basic insights in to structural and functional aspects of GlcNAc transporter mechanism, for the first time. These findings suggest that the selected residues (Asn80, Asn195 and 320 Gln320) are conserved and play pivotal roles in the protein’s structural integrity and functional mechanisms, making them prime candidates for mutagenesis studies to elucidate their precise contributions to transport activity. These results can be further extended to wet-lab experiments to test the physiological significance of critical amino acids identified using in-silico tools. This study provides strategic work flow for designing and developing effective therapeutic agents to inhibit Ngt1 activity to stop the GlcNAc entry and there by GlcNAc signaling and virulence in this commensal yeast. Thus, in-silico structural analysis can fill the gaps in wet-lab based structural determination by predicting the structure functionalities to reduce the laboratory time taking process

## Supporting information

https://www.akayhelp.in/OI2IeTJ9hRwKD30/file

## ACKNOWLEDGEMENTS

The author thanks Department of Science and Technology/SERB (CRG/2022/008507), Government of India for providing financial support. KHR acknowledges a GITAM: Research Seed Grant (RSG), Gandhi Institute of Technology and Management (GITAM), Visakhapatnam, India. The author thank the members of his lab at GITAM and members of Dr Swagata Ghosh lab at the University of Kalyani, India, for their critical reading and helpful suggestions.

The authors confirm their contribution to the paper: study conception and design: KHR; data collection: SH and NSTG; analysis and interpretation of results: SH, NSTG, SP and KHR; draft manuscript: SG, SS and KHR. All authors reviewed the results and approved the final version of the manuscript

## REFERENCES

1) Perlroth, J., Choi, B., & Spellberg, B. (2007). Nosocomial fungal infections: epidemiology, diagnosis, and treatment. Medical Mycology, 45(4), 321–346. 10.1080/13693780701218689.

2) Min, K., Naseem, S., & Konopka, J. B. (2019). N-Acetylglucosamine Regulates Morphogenesis and Virulence Pathways in Fungi. Journal of Fungi, 6(1), 8. 10.3390/jof6010008.

3) Jarvis, W. R. (1995). Epidemiology of Nosocomial Fungal Infections, with Emphasis on Candida Species. Clinical Infectious Diseases, 20(6), 1526–1530. 10.1093/clinids/20.6.1526.

4) Sullivan, D. J., Moran, G. P., & Coleman, D. C. (2005). Candida dubliniensis: Ten years on. FEMS Microbiology Letters, 253(1), 9–17. 10.1016/j.femsle.2005.09.015.

5) Cassone, A., & Cauda, R. (2012). Candida and candidiasis in HIV-infected patientspatients: where commensalism, opportunistic behavior and frank pathogenicity lose their borders. AIDS, 26(12), 1457–1472. 10.1097/qad.0b013e3283536ba8.

6) Moser, D., Biere, K., Han, B., Hoerl, M., Schelling, G., Choukér, A., & Woehrle, T. (2021). COVID-19 Impairs Immune Response to Candida albicans. Frontiers in Immunology, 12. 10.3389/fimmu.2021.640644.

7) Romani, L., Bistoni, F., & Puccetti, P. (2003). Adaptation of Candida albicans to the host environment: the role of morphogenesis in virulence and survival in mammalian hosts. Current Opinion in Microbiology, 6(4), 338–343. 10.1016/s1369-5274(03)00081-x.

8) Naseem, S., Gunasekera, A., Araya, E., & Konopka, J. B. (2011). N-Acetylglucosamine (GlcNAc) Induction of Hyphal Morphogenesis and Transcriptional Responses in Candida albicans Are Not Dependent on Its Metabolism. Journal of Biological Chemistry, 286(33), 28671–28680. 10.1074/jbc.m111.249854.

9) Noble, S. M., Gianetti, B. A., & Witchley, J. N. (2016). Candida albicans cell-type switching and functional plasticity in the mammalian host. Nature Reviews Microbiology, 15(2), 96–108. 10.1038/nrmicro.2016.157.

10) Sudbery, P. E. (2011). Growth of Candida albicans hyphae. Nature Reviews Microbiology, 9(10), 737–748. 10.1038/nrmicro2636.

11) Biswas, S., Van Dijck, P., & Datta, A. (2007). Environmental Sensing and Signal Transduction Pathways Regulating Morphopathogenic Determinants of Candida albicans. Microbiology and Molecular Biology Reviews, 71(2), 348–376. 10.1128/mmbr.00009-06.

12) Du, H., & Huang, G. (2015). Environmental pH adaption and morphological transitions in Candida albicans. Current Genetics, 62(2), 283–286. 10.1007/s00294-015-0540-8.

13) Kornitzer, D. (2019). Regulation of Candida albicans Hyphal Morphogenesis by Endogenous Signals. Journal of Fungi, 5(1), 21. 10.3390/jof5010021.

14) Alvarez, F. J., & Konopka, J. B. (2007). Identification of an*N*-Acetylglucosamine Transporter That Mediates Hyphal Induction in*Candida albicans*. Molecular Biology of the Cell, 18(3), 965–975. 10.1091/mbc.e06-10-0931.

15) Ene, I. V., Cheng, S.-C., Netea, M. G., & Brown, A. J. P. (2013). Growth of Candida albicans Cells on the Physiologically Relevant Carbon Source Lactate Affects Their Recognition and Phagocytosis by Immune Cells. Infection and Immunity, 81(1), 238–248. 10.1128/iai.01092-12.

16) Hanumantha Rao, K., Roy, K., Paul, S., & Ghosh, S. (2022). *N* LJacetylglucosamine transporter, Ngt1, undergoes sugarLJresponsive endosomal trafficking in *Candida albicans*. Molecular Microbiology, 117(2), 429–449. 10.1111/mmi.8.

17) Ghosh, S., Rao, K. H., Bhavesh, N. S., Das, G., Dwivedi, V. P., & Datta, A. (2014). *N* - Acetylglucosamine (GlcNAc)-Inducible Gene *GIG2* Is a Novel Component of GlcNAc Metabolism in Candida albicans. Eukaryotic Cell, 13(1), 66–76. 10.1128/ec.00244-13.

18) Brown, A. J. P., Budge, S., Kaloriti, D., Tillmann, A., Jacobsen, M. D., Yin, Z., Ene, I. V., Bohovych, I., Sandai, D., Kastora, S., Potrykus, J., Ballou, E. R., Childers, D. S., Shahana, S., & Leach, M. D. (2014). Stress adaptation in a pathogenic fungus. Journal of Experimental Biology, 217(1), 144–155. 10.1242/jeb.088930.

19) Su, C., Lu, Y., & Liu, H. (2016). N-acetylglucosamine sensing by a GCN5-related N-acetyltransferase induces transcription via chromatin histone acetylation in fungi. Nature Communications, 7(1). 10.1038/ncomms12916.

20) Rao, K. H., Paul, S., & Ghosh, S. (2021). N-acetylglucosamine Signaling: Transcriptional Dynamics of a Novel Sugar Sensing Cascade in a Model Pathogenic Yeast, Candida albicans. Journal of Fungi, 7(1), 65. 10.3390/jof7010065

21) Katzmann, D. J., Odorizzi, G., & Emr, S. D. (2002). Receptor downregulation and multivesicular-body sorting. Nature Reviews Molecular Cell Biology, 3(12), 893–905. 10.1038/nrm973.

22) Konopka, J. B. (2012). N-Acetylglucosamine Functions in Cell Signaling. Scientifica, 2012, 1–15. 10.6064/2012/489208.

23) Bavnhoj, L., Driller, J. H., Zuzic, L., Stange, A. D., Schiøtt, B., & Pedersen, B. P. (2023). Structure and sucrose binding mechanism of the plant SUC1 sucrose transporter. Nature Plants, 9(6), 938–950. 10.1038/s41477-023-01421-0.

24) Varadi, M., Anyango, S., Deshpande, M., Nair, S., Natassia, C., Yordanova, G., Yuan, D., Stroe, O., Wood, G., Laydon, A., Žídek, A., Green, T., Tunyasuvunakool, K., Petersen, S., Jumper, J., Clancy, E., Green, R., Vora, A., Lutfi, M., & Figurnov, M. (2021). AlphaFold Protein Structure Database: massively expanding the structural coverage of protein-sequence space with high-accuracy models. Nucleic Acids Research, 50(D1). 10.1093/nar/gkab1061.

25) Jumper, J., Evans, R., Pritzel, A., Green, T., Figurnov, M., Ronneberger, O., Tunyasuvunakool, K., Bates, R., Žídek, A., Potapenko, A., Bridgland, A., Meyer, C., Kohl, S. A. A., Ballard, A. J., Cowie, A., Romera-Paredes, B., Nikolov, S., Jain, R., Adler, J., & Back, T. (2021). Highly Accurate Protein Structure Prediction with Alphafold. Nature, 596(7873), 583–589. 10.1038/s41586-021-03819-2.

26) Tunyasuvunakool, K., Adler, J., Wu, Z., Green, T., Zielinski, M., Žídek, A., Bridgland, A., Cowie, A., Meyer, C., Laydon, A., Velankar, S., Kleywegt, G. J., Bateman, A., Evans, R., Pritzel, A., Figurnov, M., Ronneberger, O., Bates, R., Kohl, S. A. A., & Potapenko, A. (2021). Highly accurate protein structure prediction for the human proteome. Nature, 596(596), 1–9. 10.1038/s41586-021-03828-1.

27) Jumper, J., & Hassabis, D. (2022). Protein structure predictions to atomic accuracy with AlphaFold. Nature Methods, 19(1), 11–12. 10.1038/s41592-021-01362-6.

28) Morris, G. M., Huey, R., Lindstrom, W., Sanner, M. F., Belew, R. K., Goodsell, D. S., & Olson, A. J. (2009). AutoDock4 and AutoDockTools4: Automated docking with selective receptor flexibility. Journal of Computational Chemistry, 30(16), 2785–2791. 10.1002/jcc.21256.

29) Labbé, C. M., Rey, J., Lagorce, D., Vavruša, M., Becot, J., Sperandio, O., Villoutreix, B. O., Tufféry, P., & Miteva, M. A. (2015). MTiOpenScreen: a web server for structure-based virtual screening. Nucleic Acids Research, 43(W1), W448–W454. 10.1093/nar/gkv306.

30) Skrzypek, M. S., Binkley, J., Binkley, G., Miyasato, S. R., Simison, M., & Sherlock, G. (2016). TheCandidaGenome Database (CGD): incorporation of Assembly 22, systematic identifiers and visualization of high throughput sequencing data. Nucleic Acids Research, 45(D1), D592–D596. 10.1093/nar/gkw924.

31) Pao, S. S., Paulsen, I. T., & Saier, M. H. (1998). Major Facilitator Superfamily. Microbiology and Molecular Biology Reviews, 62(1), 1–34. 10.1128/mmbr.62.1.1-34.1998.

32) Abramson, J., Iwata, S., & Kaback, H. R. (2004). Lactose permease as a paradigm for membrane transport proteins (Review). Molecular Membrane Biology, 21(4), 227–236. 10.1080/09687680410001716862.

33) Kim, S., Chen, J., Cheng, T., Gindulyte, A., He, J., He, S., Li, Q., Shoemaker, B. A., Thiessen, P. A., Yu, B., Zaslavsky, L., Zhang, J., & Bolton, E. E. (2021). PubChem in 2021: new data content and improved web interfaces. Nucleic Acids Research, 49(D1), D1388–D1395. 10.1093/nar/gkaa971.

34) Wang, X., Shen, X., Qu, Y., Zhang, H., Wang, C., Yang, F., & Shen, H. (2024). Structural insights into ion selectivity and transport mechanisms of Oryza sativa HKT2;1 and HKT2;2/1 transporters. Nature Plants, 10(4), 633–644. 10.1038/s41477-024-01665-4.

35) Corringer, P.-J., Novère, N. L., & Changeux, J.-P. (2000). Nicotinic Receptors at the Amino Acid Level. Annual Review of Pharmacology and Toxicology, 40(1), 431–458. 10.1146/annurev.pharmtox.40.1.431.

36) Singh, N., Chaput, L., & Villoutreix, B. O. (2020). Virtual screening web servers: designing chemical probes and drug candidates in the cyberspace. Briefings in Bioinformatics. 10.1093/bib/bbaa034.

37) Colovos, C., & Yeates, T. O. (1993). Verification of protein structures: Patterns of nonbonded atomic interactions. Protein Science, 2(9), 1511–1519. 10.1002/pro.5560020916.

38) Carugo, O., & Djinovic-Carugo, K. (2013). Half a century of Ramachandran plots. Acta Crystallographica. Section D, Biological Crystallography, 69(Pt 8), 1333–1341. 10.1107/S090744491301158X.

39) Trott, O., & Olson, A. J. (2009). AutoDock Vina: Improving the Speed and Accuracy of Docking with a New Scoring function, Efficient optimization, and Multithreading. Journal of Computational Chemistry, 31(2). 10.1002/jcc.21334.

40) Yu, J., Shi, J., Zhang, Y., & Yu, Z. (2019). Molecular Docking and Site-Directed Mutagenesis of Dichloromethane Dehalogenase to Improve Enzyme Activity for Dichloromethane Degradation. Applied Biochemistry and Biotechnology, 190(2), 487– 505. 10.1007/s12010-019-03106-x.

41) Jones, P., Binns, D., Chang, H.-Y., Fraser, M., Li, W., McAnulla, C., McWilliam, H., Maslen, J., Mitchell, A., Nuka, G., Pesseat, S., Quinn, A. F., Sangrador-Vegas, A., Scheremetjew, M., Yong, S.-Y., Lopez, R., & Hunter, S. (2014). InterProScan 5: genome-scale protein function classification. Bioinformatics, 30(9), 1236–1240. 10.1093/bioinformatics/btu031.

42) Krogh, A., Larsson, B., von Heijne, G., & Sonnhammer, E. L. L. (2001). Predicting transmembrane protein topology with a hidden markov model: application to complete genomes11Edited by F. Cohen. Journal of Molecular Biology, 305(3), 567–580. 10.1006/jmbi.2000.4315.

43) Finn, R. D., Coggill, P., Eberhardt, R. Y., Eddy, S. R., Mistry, J., Mitchell, A. L., Potter, S. C., Punta, M., Qureshi, M., Sangrador-Vegas, A., Salazar, G. A., Tate, J., & Bateman, A. (2016). The Pfam protein families database: towards a more sustainable future. Nucleic Acids Research, 44(D1), D279–D285. 10.1093/nar/gkv1344.

44) Meng, E. C., Pettersen, E. F., Couch, G. S., Huang, C. C., & Ferrin, T. E. (2006). Tools for integrated sequence-structure analysis with UCSF Chimera. BMC Bioinformatics, 7(1), 339. 10.1186/1471-2105-7-339.

45) Yuan, S., Chan, H. C. S., & Hu, Z. (2017). Using PyMOL as a platform for computational drug design. WIREs Computational Molecular Science, 7(2). 10.1002/wcms.1298.

46) Guex, N., & Peitsch, M. C. (1997). SWISS-MODEL and the Swiss-Pdb Viewer: An environment for comparative protein modeling. Electrophoresis, 18(15), 2714–2723. 10.1002/elps.1150181505.

47) Chovancova, E., Pavelka, A., Benes, P., Strnad, O., Brezovsky, J., Kozlikova, B., Gora, A., Sustr, V., Klvana, M., Medek, P., Biedermannova, L., Sochor, J., & Damborsky, J. (2012). CAVER 3.0: A Tool for the Analysis of Transport Pathways in Dynamic Protein Structures. PLoS Computational Biology, 8(10), e1002708. 10.1371/journal.pcbi.1002708.

48) Ma, D. L., Chan, D. S. H., & Leung, C. H. (2011). Molecular docking for virtual screening of natural product databases. Chem. Sci., 2(9), 1656–1665. 10.1039/c1sc00152c

