## Supplementary material for "Expression and In-silico Structural analysis of N-acetylglucosamine transporter, Ngt1 in model fungal pathogen, *Candida albicans*": https://www.akayhelp.in/OI2IeTJ9hRwKD30/file: Supplemetary material - Copy.docx

Running title: **In-silico structural analysis of N-acetylglucosamine transporter Ngt1 in Candida**.

**Table S1. Docking energy table for Ngt1 with GlcNAc**

| **Complex** | **Mode rank** | **Lowest binding energy** | **Best mode rmsd** |
| --- | --- | --- | --- |
| **Native** | 1 | -5.4 | 0.00 |
|  | 2 | -5.3 | 27.160 |
|  | 3 | -4.8 | 28.376 |
|  | 4 | -4.6 | 28.708 |
|  | 5 | -4.6 | 29.060 |
|  | 6 | -4.5 | 28.922 |
|  | 7 | -4.3 | 29.027 |
|  | 8 | -4.2 | 21.390 |
|  | 9 | -4.2 | 26.681 |
| **Mutation at ( 80, 195, 320 )** | 1 | -5.8 | 0.00 |
|  | 2 | -5.8 | 3.731 |
|  | 3 | -5.5 | 5.467 |
|  | 4 | -5.5 | 11.643 |
|  | 5 | -5.4 | 5.678 |
|  | 6 | -5.4 | 20.863 |
|  | 7 | -5.4 | 19.965 |
|  | 8 | -5.3 | 13.946 |
|  | 9 | -5.2 | 26.067 |
| **Mutation at 80** | 1 | -5.0 | 0.00 |
|  | 2 | -5.0 | 42.708 |
|  | 3 | -4.9 | 45.320 |
|  | 4 | -4.7 | 6.461 |
|  | 5 | -4.6 | 5.272 |
|  | 6 | -4.3 | 27.555 |
|  | 7 | -4.0 | 61.025 |
|  | 8 | -4.0 | 61.709 |
|  | 9 | -4.0 | 24.485 |
| **Mutation at 195** | 1 | -6.1 | 0.00 |
|  | 2 | -5.9 | 3.226 |
|  | 3 | -5.8 | 3.379 |
|  | 4 | -5.8 | 5.405 |
|  | 5 | -5.6 | 5.580 |
|  | 6 | -5.3 | 5.366 |
|  | 7 | -5.2 | 5.416 |
|  | 8 | -5.1 | 19.048 |
|  | 9 | -5.0 | 6.585 |
| **Mutation at 320** | 1 | -5.7 | 0.00 |
|  | 2 | -5.4 | 54.434 |
|  | 3 | -5.4 | 22.981 |
|  | 4 | -5.3 | 5.619 |
|  | 5 | -5.3 | 24.101 |
|  | 6 | -5.0 | 21.955 |
|  | 7 | -4.8 | 21.605 |
|  | 8 | -4.8 | 54.931 |

**Table S2. Hydrogen bonding interactions at interaction site**

| Complex | Residue | AA | Distance H-A | Distance D-A | Donor Angle | Donor Atom | Acceptor Atom |
| --- | --- | --- | --- | --- | --- | --- | --- |
| Native | 123A | LYS | 1.84 | 2.82 | 166.87 | 1187 [N3+] | 5183 [O3] |
|  | 123A | LYS | 2.14 | 3.11 | 164.45 | 1179 [Nam] | 5173 [O3] |
|  | 124A | MET | 1.97 | 2.97 | 170.69 | 1192 [Nam] | 5178 [O3] |
| Mutation at ( 80, 195, 320 ) | 176A | SER | 2.38 | 3.14 | 134.25 | 5177 [O3] | 1706 [O2] |
|  | 247A | LYS | 1.88 | 2.80 | 152.99 | 2398 [N3+] | 5177 [O3] |
|  | 254A | MET | 2.73 | 3.50 | 134.15 | 2459 [Nam] | 5177 [O3] |
|  | 254A | MET | 2.06 | 2.76 | 127.64 | 5182 [O3] | 2462 [O2] |
| Mutation at 80 | 167A | TRP | 2.96 | 3.81 | 142.70 | 1623 [Nar] | 5173 [O3] |
|  | 289A | ASN | 1.89 | 2.85 | 158.17 | 2833 [Nam] | 5175 [O3] |
|  | 432A | LYS | 3.04 | 3.66 | 121.25 | 4423 [N3+] | 5178 [O3] |
|  | 435A | GLN | 2.12 | 2.76 | 122.12 | 5175 [O3] | 4450 [O2] |
|  | 435A | GLN | 1.97 | 2.78 | 138.86 | 5178 [O3] | 4450 [O2] |
| Mutation at 195 | 80A | ASN | 2.13 | 3.08 | 159.16 | 798 [Nam] | 5152 [O3] |
|  | 167A | TRP | 1.97 | 2.85 | 144.83 | 1622 [Nar] | 5160 [O2] |
|  | 292A | TYR | 2.18 | 3.09 | 156.49 | 5167 [O3] | 2881 [O3] |
|  | 320A | GLN | 2.22 | 2.99 | 132.61 | 3221 [Nam] | 5170 [O3] |
|  | 435A | GLN | 2.58 | 3.30 | 130.34 | 5163 [O3] | 4443 [O2] |
| Mutation at 320 | 256A | LYS | 1.84 | 2.84 | 171.92 | 2486 [N3+] | 5177 [O3] |
|  | 257A | THR | 2.63 | 3.52 | 148.83 | 2491 [Nam] | 5165 [Nox] |
|  | 257A | THR | 2.21 | 3.06 | 145.40 | 5174 [O3] | 2494 [O2] |
|  | 422A | LYS | 2.38 | 3.33 | 159.49 | 4308 [N3+] | 5177 [O3] |
|  | 491A | GLU | 1.96 | 2.76 | 137.91 | 5177 [O3] | 4987 [O2] |
|  | 492A | VAL | 2.95 | 3.58 | 123.90 | 5167 [O3] | 4997 [O2] |

**Table S3 Hydrophobic interactions at interaction site**

| Complex | Residue | AA | Distance | Ligand Atom | Protein Atom |
| --- | --- | --- | --- | --- | --- |
| Native | 122A | VAL | 3.40 | 5174 | 1177 |
|  | 123A | LYS | 3.32 | 5169 | 1185 |
|  | 242A | ILE | 3.87 | 5174 | 2337 |
| Mutation at ( 80, 195, 320) | 254A | MET | 3.80 | 5168 | 2463 |
| Mutation at 80 | 76A | PRO | 3.29 | 5169 | 760 |
|  | 80A | ALA | 3.77 | 5169 | 798 |
|  | 106A | PHE | 3.65 | 5164 | 1023 |
|  | 195A | PHE | 3.77 | 5169 | 1904 |
| Mutation at 195 | 417A | MET | 3.94 | 5163 | 4253 |
|  | 478A | ILE | 3.69 | 5163 | 4863 |
| Mutation at 320 | 106A | PHE | 3.12 | 5150 | 1022 |
|  | 432A | LYS | 3.50 | 5150 | 4404 |

**Table S4 : Parameters and their values used in the Tunnel analysis**

| Tunnel ID | Bottleneck | Length | Curvature | Throughput |
| --- | --- | --- | --- | --- |
| 1 | 2.5 | 11.8 | 1.3 | 0.84 |
| 2 | 2.5 | 15.4 | 1.5 | 0.81 |
| 3 | 2.1 | 12.7 | 1.2 | 0.81 |
| 4 | 1.9 | 15.1 | 1.3 | 0.78 |
| 5 | 1.6 | 20.1 | 1.3 | 0.70 |
| 6 | 1.4 | 16.8 | 1.5 | 0.69 |
| 7 | 1.3 | 19.9 | 1.4 | 0.60 |
| 8 | 1.4 | 16.2 | 1.3 | 0.58 |
| 9 | 1.3 | 24.7 | 1.7 | 0.52 |
| 10 | 1.3 | 38.0 | 2.1 | 0.22 |
| 11 | 0.9 | 43.1 | 2.9 | 0.10 |
| 12 | 0.9 | 64.3 | 1.8 | 0.07 |
| 13 | 1.0 | 66.2 | 2.4 | 0.04 |

**Table S5. Description of Blocker (ZINC00005512974) complex parameters**

| ZINC000005512974 | Residue | AA | | Distance H-A | | Distance D-A | Donor Angle | | Donor Atom | | Acceptor Atom |
| --- | --- | --- | --- | --- | --- | --- | --- | --- | --- | --- | --- |
| Hydrogen bonds | 170A | GLN | | 2.14 | | 3.10 | 166.29 | | 1317 [Nam] | | 29 [O3] |
|  | 411A | ILE | | 2.45 | | 3.41 | 157.95 | | 12 [N3] | | 3264 [O2] |
|  | 424A | THR | | 2.55 | | 3.40 | 145.78 | | 3367 [O3] | | 22 [O3] |
|  | Residue | | AA | | Distance | | | Ligand Atom | | Protein Atom | |
| Hydrophobic Interactions | 174A | | ILE | | 3.22 | | | 5 | | 1344 | |
|  | 183A | | LYS | | 3.36 | | | 19 | | 1417 | |
|  | 424A | | THR | | 3.13 | | | 24 | | 3366 | |
|  | 428A | | ALA | | 3.15 | | | 10 | | 3396 | |

| PROPERTIES | ZINC000005512974 (Brucine) |
| --- | --- |
| Lipinski Rule | Yes; 0 violation |
| Ghose | Yes |
| Veber | Yes |
| Egan | Yes |
| Muegge | Yes |
| Bioavailability score | 0.55 |

**Table S6. Virtual Screening top table:**

| Compound | IUPAC names | Lowest binding energy | structures |
| --- | --- | --- | --- |
| MolPort-001-742-110_ZINC000043552589 | 8-[5-[(2S)-5,7-dihydroxy-4-oxo-2,3-dihydrochromen-2-yl]-2-hydroxyphenyl]-5,7-dihydroxy-2-(4-hydroxyphenyl)chromen-4-one | -11.5 | 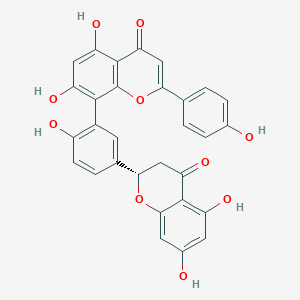 |
| MolPort-039-052-621_ZINC000044404209 | (2S)-2-[4-[5-[(2S)-5,7-dihydroxy-4-oxo-2,3-dihydrochromen-2-yl]-2-hydroxyphenoxy]phenyl]-5,7-dihydroxy-2,3-dihydrochromen-4-one | -11.3 | 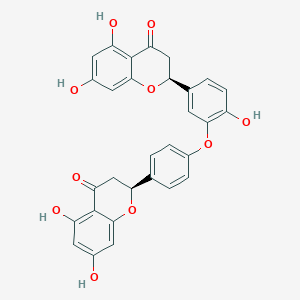 |
| MolPort-000-704-745_ZINC000005512974 | [**Brucine**](https://zinc12.docking.org/synonym/Brucine)([Brucine:Strychnidin*-*10-one,2,3-dimethox*y*](https://zinc12.docking.org/synonym/Brucine:Strychnidin-10-one,2,3-dimethoxy)) | -10.1 | 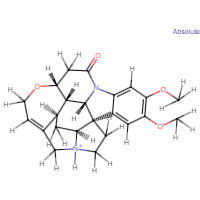 |

**Supplementary Text**

ZINC000005512974: The analyzed molecule, ZINC000005512974 provided, exhibits a range of physicochemical, pharmacokinetic, drug-likeness, and medicinal chemistry properties that highlight its potential for drug development. The molecular formula C23H27N2O4+ corresponds to a molecular weight of 395.47 g/mol, with 29 heavy atoms and 6 aromatic heavy atoms, while the fraction of carbon atoms in an sp3 hybridization state is 0.61, reflecting a moderate level of saturation. The molecule has 2 rotatable bonds, 4 hydrogen bond acceptors, and 1 hydrogen bond donor, alongside a molar refractivity of 115.00 and a topological polar surface area (TPSA) of 52.44 Å², which together influence its potential interactions and bioavailability[ Lipinski et al., 2011].

The lipophilicity profile, assessed using multiple methods, indicates a consensus Log P value of 0.87, with individual estimations such as 3.25 (iLOGP), 0.98 (XLOGP3), -0.07 (WLOGP), 1.84 (MLOGP), and 1.84 (SILICOS-IT). This range of values suggests a balanced lipophilic character suitable for drug development(Veber et al. 2002). Solubility assessments revealed a Log S (ESOL) value of -2.93, equating to a solubility of 4.64 x 10^-1 mg/ml, classifying it as soluble. Additional solubility predictions, such as Log S (Ali) at -1.67 and Log S (SILICOS-IT) at -3.61, indicate varied but consistent solubility properties, with classifications spanning soluble to very soluble[ Egan et al., 2000].

Pharmacokinetically, the molecule is characterized by high gastrointestinal absorption but does not permeate the blood-brain barrier (BBB). It is not a substrate for P-glycoprotein (P-gp) and does not inhibit major cytochrome P450 enzymes (CYP1A2, CYP2C19, CYP2C9, CYP2D6, and CYP3A4), indicating a reduced risk for drug-drug interactions mediated by these enzymes[ **Gleeson, 2008**]. The skin permeability, represented by a Log Kp of -8.02 cm/s, suggests limited transdermal diffusion.

Regarding drug-likeness, the molecule adheres to Lipinski’s rule of five without any violations, enhancing its oral bioavailability profile[ Lipinski, 2016]. It also passes other key drug-likeness filters, including the Ghose, Veber, Egan, and Muegge criteria, further supporting its favorable properties for drug development[ Jia et al., 2020]. The bioavailability score of 0.55 underscores a moderate likelihood of achieving effective oral absorption(Brenk et al. 2008a).

In terms of medicinal chemistry, the molecule raises no Pan Assay Interference Compounds (PAINS) alerts(Anon n.d.-c), indicating a lack of structural features associated with non-specific binding. However, one Brenk alert for an isolated alkene is noted, which could be considered in further optimization[ Brenk et al., 2008]. Despite a molecular weight above 350 g/mol, rendering it non-leadlike by Leadlikeness criteria, its synthetic accessibility score of 5.48 reflects moderate synthetic feasibility.

This comprehensive analysis demonstrates the molecule’s balanced physicochemical and pharmacokinetic attributes, favorable drug-likeness, and manageable medicinal chemistry considerations, making it a promising candidate for further optimization in drug discovery efforts.
